# Canonical G-protein coupled receptors of vascular plants

**DOI:** 10.64898/2026.03.02.708220

**Authors:** Valentina Fernandez-Figueroa, Camila A. Quercia-Raty, Jan Gallastegui-Ulloa, Luka Robeson, Sebastian E. Brauchi

## Abstract

G-protein coupled receptors (GPCRs) are responsible for translating environmental signals of various types into cellular signals. Over 40 thousand GPCR orthologs have been discovered in the supergroup Unikonta, and around 800 genes encode for GPCRs in the human genome. In contrast to this astonishing variety, only a handful of GPCR-related genes have been reported in vascular plants, a major group within land plants. In an attempt to advance our understanding of plant GPCRs as well as their role in plant cellular signaling, here we present comprehensive bioinformatic analysis that includes phylogenetic hypotheses, *in silico* structural analysis, and tissue distribution of transcripts. Altogether, our work strongly suggests that GCR1 is the sole genuine GPCR expressed in Embriophyta. Finally, we briefly discuss the potential role of GCR1 in root hairs, the tubular outgrowths in root epidermal cells that are involved in nutrient absorption, environmental interaction, and root development.

## INTRODUCTION

The survival of living organisms relies on the ability to detect, integrate, and respond to environmental signals. The G Protein Coupled Receptor (GPCR) superfamily plays a fundamental role in the transduction of environmental cues into coherent cellular signals.

GPCRs are membrane protein receptors, displaying 7 transmembrane helices (7TM). Thought to be descendants of bacterial rhodopsins (Taylor & Agarwal, 1993) their expression has been reported at all levels of the tree of life, from unicellular eukaryotes such as *Saccharomyces cerevisiae* (Wang et al., 2021)*, fungi* (Gruber et al., 2013) social amoebas such as *Dictyostelium discoideum* (Raisley et al., 2004), and single celled green algae (Port et al., 2013), to more complex organisms such as vertebrates (Sakai et al., 2019) and land plants (Amora et al., 2016). Only in humans, more than 800 individual genes are known to encode for unique GPCR proteins (Stevens et al., 2013). In contrast to the vast literature on GPCR function in Metazoa, there is scarce information and controversy regarding the presence, number, and function of GPCR proteins in vascular plants (Chakraborty & Raghuram, 2023).

### GPCR mechanism of transduction

GPCR signalling diversity arises from a multilayer detection architecture, combining receptor multiplicity, selective coupling to distinct Galpha subunit and transactivation mechanisms that expand signaling output beyond ligand-receptor interaction (Masuho et al., 2023).

An additional layer of complexity emerged from combinatorial assembly of heterotrimeric G protein subunit engaged for individual GPCRs, enabling signal diversification downstream of receptor activation (Eichel, Kelsie and von Zastrow, 2018; Masuho et al., 2021). For example in humans, this combinatorial potential of GPCR signaling machinery is supported by 33 G protein-encoding genes, comprising 16 Galpha, 5 Gbeta, and 12 Ggamma subunits (Hillenbrand et al., 2015; Yang & Hildebrandt, 2006).

In general, Galpha subunits are classified into four functional families: Gs, Gi/o, Gq/11, and G12/13 (Neves et al., 2002). Furthermore, Galpha activity is modulated through interactions with proteins such as Regulators of GPCR Signaling (RGS). These RGS proteins possess domains that act as GTPase-activating proteins (GAPs) or guanine dissociation inhibitors (GDIs) allowing precise control over the GTPase cycle of Galpha subunits (Bradford et al., 2013; De Mendoza et al., 2014).

In contrast to Metazoans, vascular plants display a significantly reduced number of G protein-encoding genes: a sole canonical Galpha subunit (GPA1), only one Gbeta subunit (AB1), and three Ggamma (AGG1, AGG2A, and AGG3) (Urano et al., 2013). Additionally, three extra-large Galpha proteins (XLGs) have been reported in plants (Maruta et al., 2021).

While this streamlined repertoire of molecular transducers limits the combinatorial possibilities for GPCR signaling in vascular plants compared to metazoans, it presents an intriguing paradox: despite lacking the capacity for rapid sensory adaptation through a diverse set of GPCRs, plants seems to utilize G protein-mediated signaling pathways efficiently (Mohanasundaram & Pandey, 2023). Increasing evidence confirms G protein involvement in critical processes dealing with fundamental plant physiology (T. Y. Wu et al., 2022). Nevertheless, fundamental questions persist regarding the precise interplay between putative plant GPCRs, their cognate G proteins, downstream effectors, and secondary messengers, a knowledge gap that underscores the need for deeper mechanistic investigation.

### GPCR classification

According to the GRAFS classification system (Dale et al., 2022; Krishnan et al., 2012), GPCRs can be divided into 6 classes: Class A (Rhodopsin), Class B (Secretin and Adhesion), Class C (Glutamate), Class D (Fungal pheromone receptor), Class E (cAMP receptor), and Class F (Frizzled/Taste2) (Eilers et al., 2005; Fredriksson, Höglund, et al., 2003; Fredriksson & Schiöth, 2005; Schiöth & Fredriksson, 2005). This classification is based on their sequence and function, where each family has a unique set of signature residues that allows grouping (Do et al., 2022; Isberg et al., 2015).

While classes A, B, and C contain representatives widely distributed in Metazoa and a handful of unicellular eukaryotes (Kooistra et al., 2021), classes D and E are almost exclusively found in fungi and amoeba, respectively (Z. Liu et al., 2025). Finally, class F GPCRs appear widely distributed throughout Unikonta (Krishnan et al., 2012).

Although the GPCR diversity in Unikonta is vast, a striking contrast is observed in the Viridiplantae (the large group of green plants) where only a handful of candidates have been proposed to be GPCRs expressed in vascular plants such as *Nicotiana tabacum* (Apone et al., 2003), *Oryza sativa* (Ma et al., 2015), Solanum lycopersicum (Amora et al., 2016), Upland cotton (Lu et al., 2018) and *Arabidopsis thaliana* (Pandey et al., 2009).

Overall, the recent efforts to report and characterize GPCR-type proteins in plants have identified a limited set of gene products including GCR1, GTG/COLD 1 and 2, CAND1, CAND3, CAND 8, and CAND 9 (Gookin et al., 2008).

Aiming to provide a deeper insight into the GPCR repertoire in plants, we combined phylogenetic reconstructions and structural clustering to uncover bona fide GPCR-encoding genes. Our analysis revealed that GCR1 is the sole canonical GPCR encoded in vascular plants. Moreover, we found that GCR1 contains a combination of structural motifs associated with class A, B and E families. Finally, we surveyed tissue expression indicating that the protein is highly expressed in root hairs, an extension of plant epidermal cells operating as a sensory component, characteristic of the root epithelia.

## RESULTS

### Identifying and selecting putative GPCR proteins in *Arabidopsis thaliana*

Aiming to make a comprehensive protein sequence dataset, we first conducted a screening of plant GPCR protein candidates in *A. thaliana*. To this end, we took into account the few proposed GPCR-like genes, collecting an initial set of 22 candidate proteins that are present in *A. thaliana* (Supplementary Table 1). Aiming for the canonical fold, we selected at least one protein per group, prioritizing those with 7 predicted transmembrane domains, a keystone characteristic structural feature of GPCRs, making a final set of 17 candidates. The set includes G-protein coupled receptor 1 (GCR1)(Plakidou-Dymock et al., 1998), *Mildew Locus O* (MLO) proteins 1 and 5 (Chen et al., 2009; Iovieno et al., 2015), Tobamovirus Multiplication proteins TOM1 (Lu et al., 2018) the Heptahelical transmembrane protein 2 (HHP2) (Hsieh & Goodman, 2005), and 8 proteins classified as *candidate* (CAND 1-5, 8-10) (Gookin et al., 2008).

### Clustering of putative plant GPCRs in the context of current GPCR classification

Once we collected the curated list of putative plant GPCRs, we performed a phylogenetic reconstruction to infer classifications according to the generally accepted GPCR categorization. For this purpose, we included well characterized members from all six GPCR families according to the current GRAFS classification giving preference to proteins with an experimentally resolved structure according to GPCRdb (https://gpcrdb.org/) (Supplementary Table 1). Also, we included sequences from multiple model organisms (including members from metazoa, fungi and protista) to maximize representativeness and support a reliable classification of plant GPCR candidates within GPCR diversity. Thus, the final set comprises 29 non-redundant sequences from Unikonta (retrieved from the GPCRdb and GPCR PEnDB), together with the 17 putative GPCR sequences from *A. thaliana*, previously defined.

Following the MSA derived from our curated database, we clustered the primary sequences according to a pairwise distance matrix in a UPGMA tree (Figure 1A). Our analysis showed that the full set of GPCR sequences are recovered in one large clade containing all members from Class A, B, C, D, E and F (GPCR Clade, Figure 1A), except by the human orphan receptor RAI3 (member of class C). All plant GPCR candidates appeared outside of the GPCR clade, with GCR1 being the sole notable exception. GCR1 is contained in a clade comprising type E and B GPCR. Type E GPCR are found in social amoebas (Amoebozoa), sister group to Opisthokonta (Kawabe et al., 2020).

**Figure 1:**
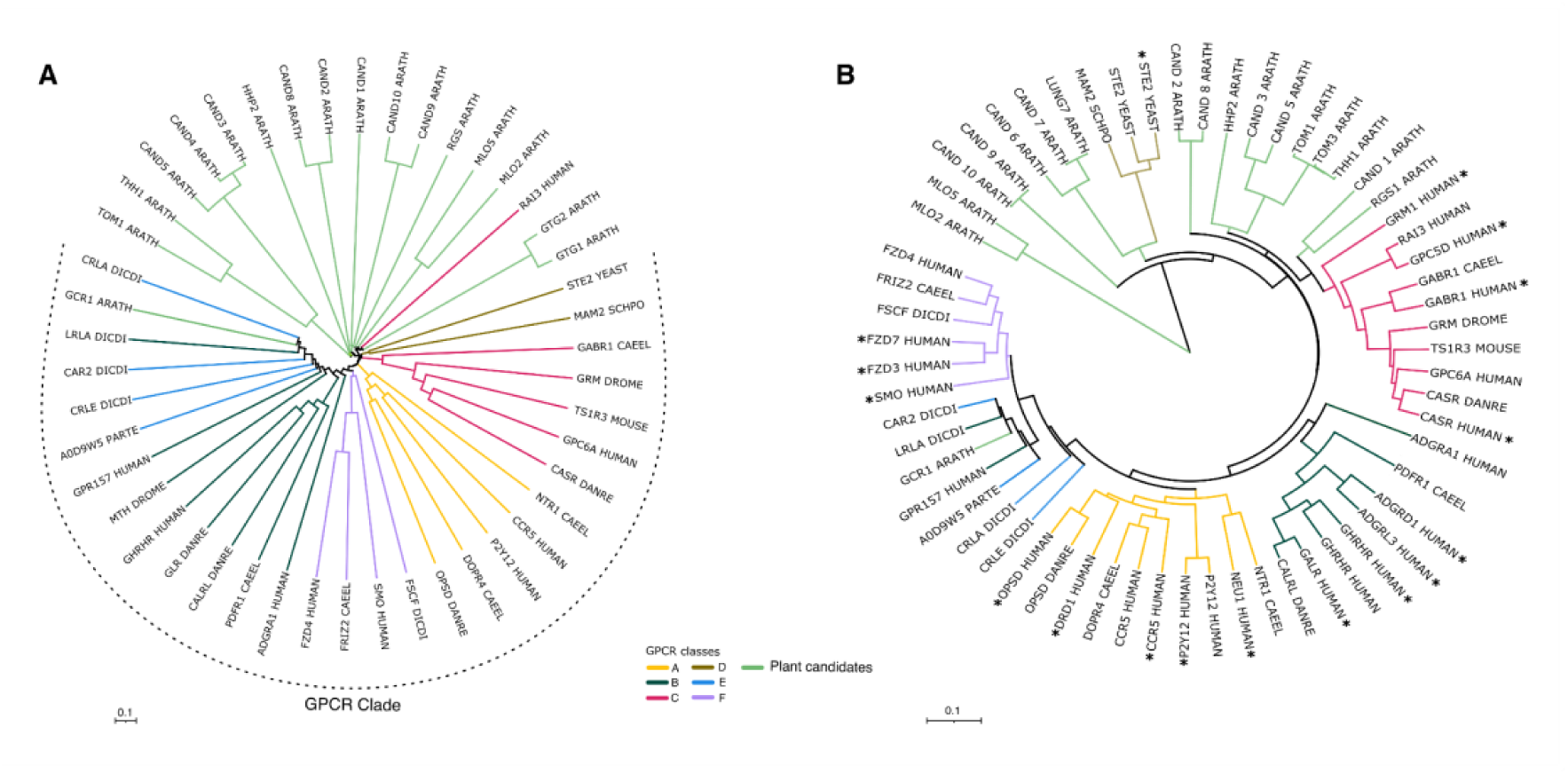
Phylogenetic and structural clustering of Arabidopsis GPCR candidates within the GPCR superfamily. **A)** UPGMA tree obtained from protein primary sequence alignment. Tree scales show distinct site proportion (p-distance) **B)** UPGMA tree displaying distances among protein structures alignment considering only transmembrane structures. Tree scales show structural dissimilarity (1 − TM-score). Asterisks indicate experimentally resolved protein structure. Plants GPCR candidates are shown as green branches, while bona fide GPCR are grouped according to classes: Class A (yellow), B (dark green), C (pink), D (light brown), E (blue) and F (purple). Notably, GCR1 is the only plant GPCR candidate clustering within a canonical GPCR clade, grouping with members of families B and E.

Protein structure is closely related to function and sensitive to distant homologies. For this reason, we decided to search for structural similarities to further confirm the position of GCR1 in the context of canonical GPCRs. To this end, a pairwise structural alignment with the TM-align algorithm was performed between the transmembrane domains of predicted AlphaFold models of our gene set and experimental structures of canonical GPCRs (Supplementary Table 1). The resulting pairwise distance matrix was clustered using UPGMA and represented as a dendrogram (Figure 1B) Note that the most distant outliers detected in our initial phylogenetic analysis (i.e. AtGTG1 and AtGTG2 with 9 and 10 TM respectively) were not further considered. Consistent with the accepted GPCR categorization (Fredriksson, Lagerström, et al., 2003), the obtained dendrogram from TM-align shows a monophyletic clades for Class C, D and F as well as a polyphyletic clade containing sequences from class A, B and E families (Figure 1B). This result is in agreement with previous hypotheses of GPCR evolution (Krishnan et al., 2012). As expected MLO proteins are located at the root of the tree, not related with the clades defined by bona fide GPCRs.

GPCR sequences as well as *A. thaliana* candidates appear well grouped by receptor sub classification. The only exception is class D GPCRs, nested as sister to CAND 6-7 candidates. GCR1, appear well nested as part of the clade formed by sequences of class B and E. All the remaining putative proteins (i.e., CAND, TOM, and HHP2) group separately from robust GPCR groups (Figure 1B, green branches) and therefore considered non-GPCR related.

Within the clade containing GCR1 we found the human ortholog GPR157, a receptor expressed in radial glial progenitor cells (RGP) (Takeo et al., 2016), confirming a recent hypothesis (Gotkhindikar & Chakravorty, 2025). Additionally, a class E GPCR from *Dictyostelium discoideum* (Uniprot id= P34907) is located sister to GCR1 (Figure 1A and 1B).

### GCR1 orthology and conservation

Thus, according to our analyses, GCR1 would likely be the only canonical GPCR protein present in *A. thaliana*’s genome. Consequently, we explored the orthology of GCR1 amongst Eukarya, including Metazoa, Unicellular heterotrophs (protists) and Viridiplantae to address gene robustness over a large evolutionary time span (Figure 2). To this end, we collected available coding nucleotide sequences, corresponding to one-to-one and one-to-many orthologs of AtGCR1 (Supplementary Table 2).

**Figure 2:**
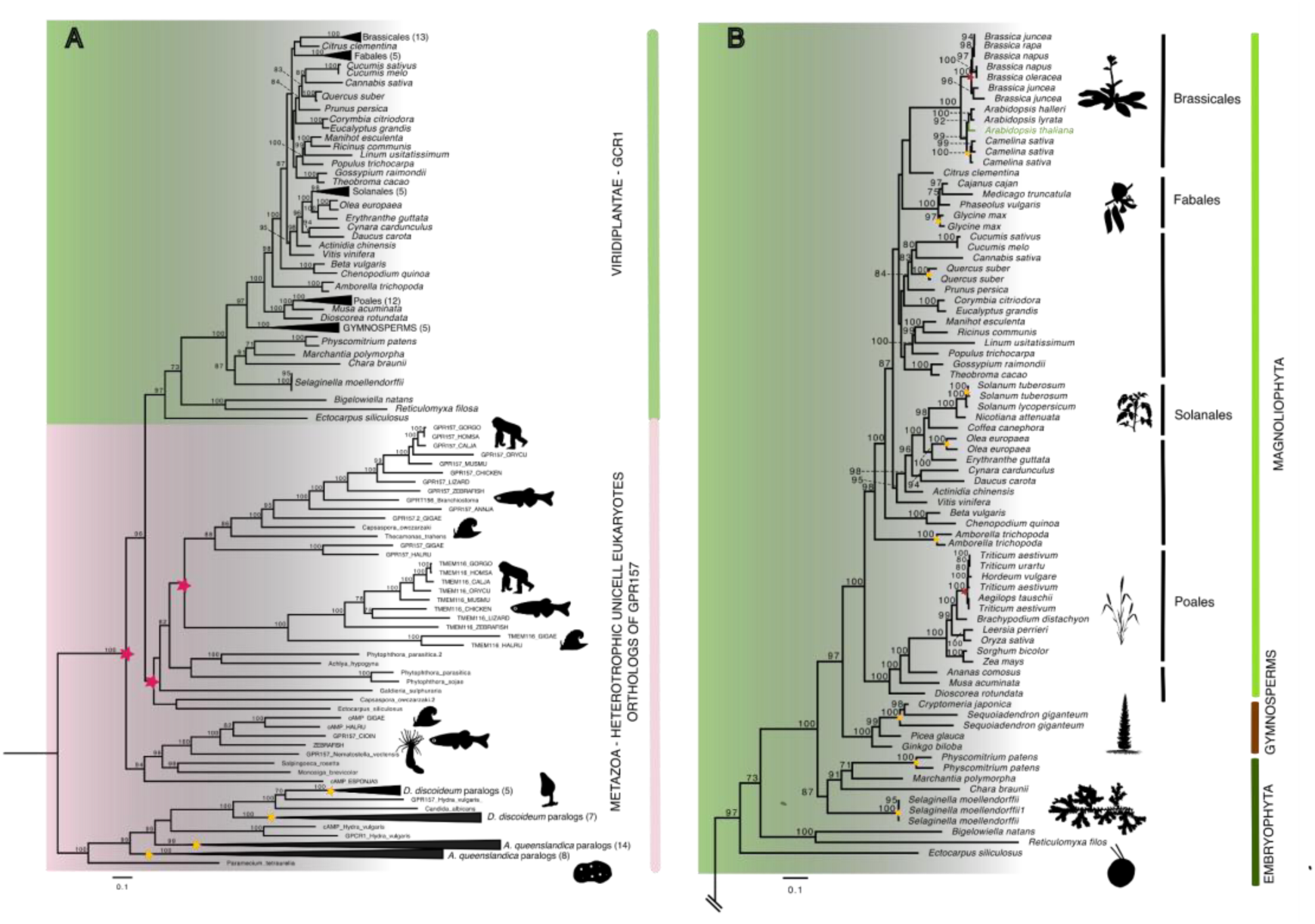
Phylogenetic Maximum Likelihood tree, displaying the evolutionary relationships of GCR1 gene across model Eukaryotes species. The tree was rooted at midpoint. The number above the nodes indicate bootstrap support values. **A)** Expanded tree including metazoan and unicellular heterotrophs (light pink highlight) and plant orthologs (green highlight). Stars denote nodes with duplication events (red for ancestral duplications, yellow for intraspecific duplications). In collapsed clades, parenthesis indicate the number of sequences per clade. **B)** Inset of Plants orthologs tree for AtGCR1. Stars denote duplication events. (red for ancestral duplications, yellow for intraspecific duplications). The green branch indicates AtGCR1. Black vertical lines indicate taxonomic orders and colors vertical lines indicate taxonomic divisions.

With a dataset of 157 unique CDS, we next performed a Maximum likelihood Phylogenetic analysis (Figure 2), to describe the evolutive dynamics (Figure 2A) and diversification of the gene across land plants (Figure 2B). Our phylogenetic reconstruction unveiled an intricate evolutionary history of the GCR1 gene, where the evolutionary history of the gene does not match the evolutionary history of the species in invertebrates and unicellular heterotrophs. Ancients duplications (Figure 2A; red stars) suggests a birth-and-death gene evolution (Nei & Rooney, 2005), for instance, the social amoeba *D. discoideum* shows 12 copies, Choanoflagellates *(S. rosetta* and *M. brevicolor*) shows only one copy and the sponge *A. queenslandica* display 23 copies, while vertebrates has only two copies of the gene (GPR157 and TMEM116).

In contrast, AtGCR1 orthologs are well conserved in Viridiplantae. Most species exhibit a single copy gene, some exceptions are lineages with whole genome duplication as *Brassicas, Cameliana sativa, Glycine max, Solanum tuberosum, Triticum aestivum, Physcomitrium patens*. This observation suggests a tight control of expression and purifying selection in plants. Interestingly, some species such as *Quercus suber, Olea europea, Amborella trichopoda and Selaginella moellendorffii* present more than one copy (Figure 1A; yellow stars). All these observations reinforce the proposed birth-and-death gene evolution hypothesis.

### Sequence analysis and structural predictions for AtGCR1

Structurally, GPCRs are membrane proteins containing seven transmembrane (TM) helices arranged presenting different ligand-binding sites in their ectodomain and at the center of the transmembrane region (D. Zhang et al., 2015). Importantly, the different GPCR subgroups display a series of non-contiguous amino acids defining fingerprint motifs that are characteristic of each GPCR family. Thus, we decided to explore the signature elements present in plant GCR1 orthologs as well as the signature elements present in AtGCR1 specifically.

The MSA corresponding to all GCR1 orthologs collected,used to construct the sister relation between them, showed that GCR1 is well conserved in plants, sharing over 60% of identity within viridiplantae, and 41.9% of identity between AtGCR1 and the distant gene in *Chara braunii*.

Interestingly, the consensus sequence obtained displays 7 fingerprint motifs, corresponding to characteristic signatures that are segregated in the different metazoan GPCR families (Figure 3A).

**Figure 3.**
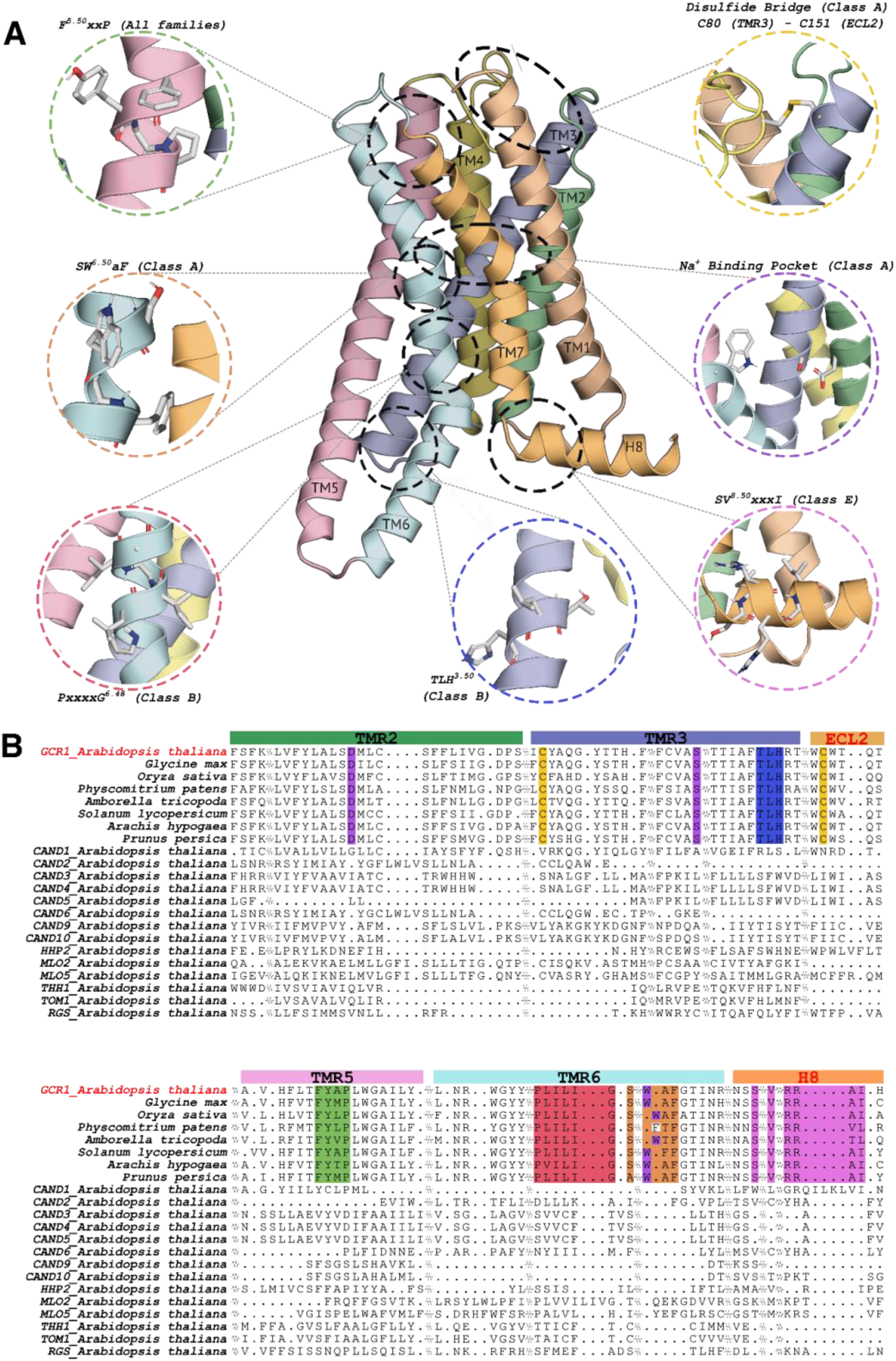
Structural characterization of AtGCR1 and conservation of GPCR-associated motifs. **A)** Predicted three-dimensional structure of AtGCR1. Insets highlight structural motifs resembling those described in metazoan GPCR families. **B)** Multiple sequence alignment showing conserved motifs (color-coded) across plant GCR1 orthologs and A thaliana candidate GPCRs identified in this study that contain seven transmembrane domains. Transmembrane segments are indicated above the alignment, with colors matching the respective transmembrane regions in the GCR1 structural model.

In agreement with the calculated distance tree, we observed that GCR1 orthologues display almost every conserved motif described in family E. These signatures include the presence of the F^5.50^xxP^5.53^ motif within TM5, a highly conserved signature across GPCR families that plays a pivotal role in helix packing and receptor conformational flexibility. Moreover, a canonical disulfide bridge between Cys80 and Cys151, linking TM3 and ECL2, would be likely preserved, suggesting a structurally stable extracellular domain. The SW^6.50^xF motif in TM6, a contraption of the CW^6.50^xP motif present in Rhodopsin-like GPCRs critical for the transition from inactive to active receptor states, is also retained in GCR1, indicating functional readiness for G-protein coupling.

Notably, GCR1 displays a partial conservation of a sodium binding pocket that is characteristic of family A members (Ben-Chaim et al., 2006). The canonical pocket in Type A GPCRs would be centered at Asp2.50 and supported by S3.39, W6.48, N7.45 and N1.50 forming an hydrogen-bond cage that would stabilize the inactivated state (Katritch et al., 2014). In GCR1, both alignment and structural prediction unveils sequence and spatial conservation of D^2.50^, S3^.39^ and W^6.48^ at the coordinating positions, suggesting a potential for analogous sodium coordination at the receptors core and the possibility of voltage-dependent amplification as part of the GCR1 response.

Remarkably, GCR1 also contains elements characteristic of class B receptors, such as the TLH^3.50^ motif and a divergent variant of the P^6.41^xxG motif, here observed as P^6.41^xxxxG^6.46^, highlighting both evolutionary conservation and adaptive divergence and suggesting a possible conserved unfold and kink in TM6 upon activation retained by TLH motif in the inactive state of the receptor.

Additional features include the SVxxxI motif, often associated with the stabilization of helix 8 in class E GPCRs at the inner membrane space (Sato et al., 2016), further underscoring the mosaic architecture of GCR1.

Collectively, the presence of these conserved structural signatures—spanning transmembrane helices and extracellular domains—strongly supports the notion that GCR1 corresponds to a canonical GPCR that was evolutionarily retained in plants. The unique combination of motifs from multiple GPCR lineages, coupled with a highly conserved topology, further suggests that GCR1 represents an ancestral scaffold, predating the diversification of GPCR families in metazoa.

### Tissue expression of GPCR-like genes in vascular plants

To elaborate further on the possible physiological roles of GCR1 in plants, we inspected the available tissue-specific transcriptomic data. The used resources integrate large-scale RNA-seq datasets covering multiple cell types, developmental stages, and anatomical structures in Arabidopsis thaliana (see methods). Overall, our analysis revealed that GCR1 expression is predominantly localized to root tissues (Figure 4). Transcripts were highly enriched in specific zones of the root system. Both a genome-wide analysis as well as high-resolution developmental transcriptome (Klepikova et al., 2016) revealed a pronounced signal in radicle during embryogenesis and signal in elongation and maturation zones of the root, supporting a role for GCR1 in cell type specific signaling processes during root development and differentiation (figure 4A and 4B).

**Figure 4.**
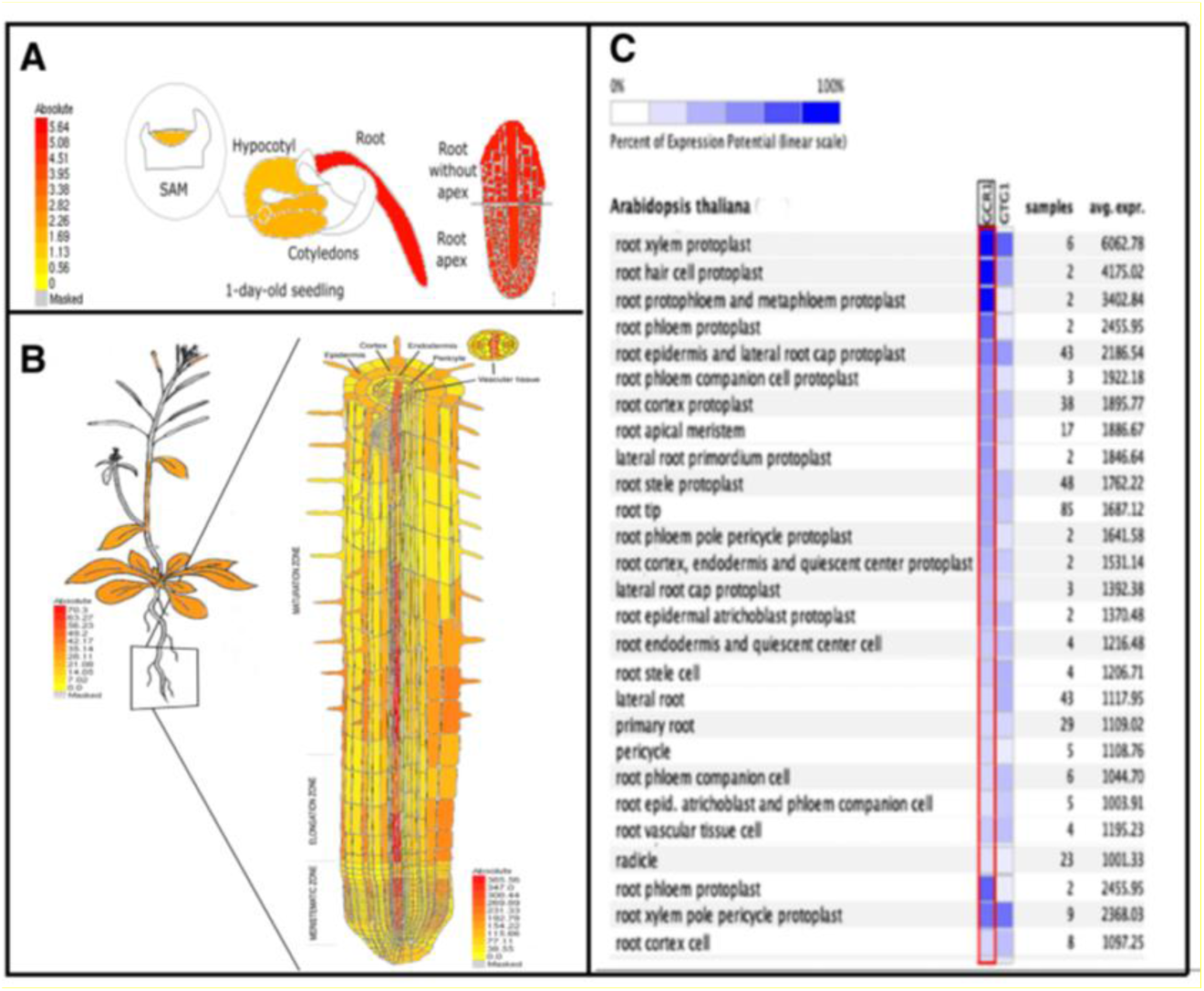
Multi-scale expression landscape of Arabidopsis thaliana GCR1. **A)** Tissue-level expression map derived from the Klepikova developmental atlas showing the preferential GCR1 expression in underground organs. **B**) Expression profile along a mature primary root revealing epidermal expression restricted to root hair cells. **C**) Single cell heatmap of isolated root cells population revealing high expression in root hairs cells, vascular - associated cells and xylem pole pericycle.

The analysis revealed that GCR1 transcripts are highly enriched in root tissues, particularly in root xylem protoplasts, root hair cells and protophloem/metaphloem protoplast, indicating a preferential expression in vascular-associated and specialized epidermal cells (figure 4C). These findings suggests a the functional relevance of GCR1 in distinct root cell populations.

## Discussion

A number of GPCR candidates have emerged in plant research over the years. Here, armed with current informatic tools, we decided to revisit these analyses. First, both our clustering analysis and structural classification places AtGCR1 within canonical GPCRs. Our analysis firmly positions GCR1 as the sole GPCR in *Arabidopsis thaliana* and other vascular plants. The absence of a validated ligand or mechanism of activation places GCR1 under the formal category of orphan GPCR.

Additionally, our findings present a false paradox where a variety of functional heterotrimeric G protein machinery is involved in a vast array of critical processes, yet plants display just one GPCR *antenna* to interact with. In contrast, the variety of inputs detected by metazoan GPCRs makes it easier to grasp whether a cell detects an external cue of different nature. That seems absent or at least broken in plant cells. That premise might be somewhat false because it assumes that a canonical GPCR would be the sole responsible for G protein-associated signaling, while in reality, the interaction of G proteins with several receptors from Viridiplantae have been demonstrated. One example would be the leucine-rich repeat (LRR) receptor-like protein FEA2 from maize, an ortholog of *Arabidopsis* CLAVATA1 that interacts with GPA1 mediating immune responses through pattern-triggered immunity (PTI) (Ishida et al., 2016). Another example are members of the MLO (mildew resistance locus O) family, which possess 7TM domains but do not share the canonical GPCR scaffold or motifs, as demonstrated here; yet they have been shown to engage with G protein subunits in signaling contexts (Lorek et al., 2016).

Second, previously studied GCR1 orthologs from vertebrates (including human), amoebozoa (*D. discoideum*), and Paramecium, underscore a putative conserved role in coupling environmental cues to cell differentiation and growth control (Takeo et al., 2016; Van Houten et al., 2000; L. Wu et al., 1995).

In this context, GCR1 emerges not as the sole G proteins interactor in plants, but rather as the only one that fits the structural and evolutionary definition of a true GPCR. This suggests that plants have evolved a unique strategy involving a combination of a single GPCR dedicated to modulate signaling pathways, alongside a variety of non-canonical receptor-like proteins that acquire the ability to couple to G proteins to signal otherwise.

### The potential role of GCR1 in root hairs

The highly specific expression in root hairs of AtGCR1, suggests a role in environmental sensing at the root-soil interface and remains to be experimentally tested.

Interestingly, our comparative sequence analysis revealed that GCR1 retains three out of the five critical residues forming the Na+ binding pocket, a feature displayed by Type A GPCR members. In Type A GPCRs, the presence of sodium on this site is essential for stabilizing the inactive state via a voltage-dependent mechanism (W. Liu et al., 2013; Venkatakrishnan et al., 2013). It is tempting to hypothesize that the highly negative membrane potential of root hairs (−120 to - 200mV) could favor Na^+^ binding, somewhat compensating for the absence of two canonical residues at the binding pocket. At the same time, a voltage-dependent mechanism would favor receptor stabilization in the inactive state under membrane resting conditions, and might represent an adaptation to the plant cellular context. This might be important considering that GPA1 in arabidopsis is constitutively active, then GCR1 stabilized in the inactive state would sequester Gɑ, effectively inhibiting downstream signalling. We can envision that GPCR voltage dependence can link local depolarization coming from, for example, OSCA channels triggered by mechanical activation into a signaling cascade, without the need of any ligand. In this scenario, a sudden depolarization would relieve Gɑ, activating the signalling cascade, amplifying the mechanical response occurring when roots explore the soil.

Under the light of our bioinformatics, we suggest that GCR1 might represent an ancient version of GPCRs. Thus, it would be a significant leap forward in the GPCR field to clarify what exactly activates GCR1, as well as directly testing its downstream signaling. These might provide a deeper understanding of the receptor mechanics and have the potential to provide a missing link to the ancestral mechanism of G protein activation.

## METHODS

### Collection and UPGMA analyses of GPCR plant candidates and canonical GPCR from other organisms

We collected primary protein sequences of the GPCR plant candidates previously described in the literature, such as CAND, TOM, HHP2, THH1, GCR1, GTG, MLO and atRGS, from uniprot databases (see Supplementary Table 1 for references and protein details), thus obtaining a data set with 17 putative plant GPCRs. Next, we searched for bona fide metazoan GPCRs of each GPCR class (A-E). For this purpose we looked into GPCRs databases (https://gpcrdb.org/) and GPCR PEnDB (https://gpcr.utep.edu/database) (Begum et al., 2020) and gave priority to proteins with experimental structures. This last was possible for most proteins, except class E (cAMP receptors) and class D (Fungal mating pheromone receptors), which are specific from amoebozoa and fungi organisms, respectively. Multiple model species were included for this data collection in order to reduce possible bias due to the distant evolutive relationship between *A. thaliana* and *Homo sapiens*. In total, we obtained 29 GPCR bona fide primary sequences, making a final dataset with 46 protein sequences. After calculation of the number of transmembrane segments with TMHMM (Hallgren et al., 2022), we perform a MSA on MAFFT website allowing the software to choose the strategy to determine best fit. An UPGMA dendogram reconstruction was made on the same website, visualized on figtree V.1.4.4 (Rambaut et al., 2018) and prepared for image publication on Inkscape V1.4.2.

We recovered the modelled structure for all the sequences in our list from the AlphaFold Database (Jumper et al., 2021). Experimental structures of canonical GPCRs from families **A** (PDB ID: 7F1R, 7X2F, 7QVM, 7XXI, 5W0P), **B** (7SF7, 7TYO, 7CZ5, 7WU2), **C** (9ASB, 7EB2, 7DGE, 9IMA), **D** (7AD3) & **F** (8YY8, 6XBM, 8JHI) were retrieved from the Protein Data Bank (Berman, 2000). No experimental structures of members of family E could be found at this time. Structures were trimmed of their N and C termini, and any long intra- and extracellular loops. The remaining transmembrane domains were aligned using TM-Align (Y. Zhang & Skolnick, 2005) and the resulting pairwise TM scores were used as similarity scores for a subsequent UPGMA clustering.

### Phylogenetic analyses of GCR1 orthologs across Eukarya

To collect GCR1 orthologs and paralogs, we took into account the information displayed by Ensembl (Dyer et al., 2025), PhyloGen at The Arabidopsis Information Resource (TAIR) on www.arabidopsis.org, and PLAZA dic 5.0 (Van Bel et al., 2022) databases. For gymnosperm and green algae species, we look forward to BLAST on the NCBI and Ensembl platform. Thus, we obtain the CDS sequences from major groups of vertebrates, metazoan model species, major groups of Viridiplantae and unicellular eukaryotic organisms. After we corroborated the accuracy of CDS translation and the number of transmembrane segments with TMHMM (Hallgren et al., 2022) we obtain a final data set with 157 number of sequences (See details of the taxonomy sampling and the accession numbers on Supplementary Table 2). Some instances appear with nucleotide sequences incomplete; we took these sequences in our analyses, because the different incomplete sequences of one species appear to complete the canonical one (i.e. *Sequoiadendron giganteum*). Multiple Sequence Alignment (MSA) was performed using MAFFT (Katoh et al., 2018; Katoh & Standley, 2013) on its web version, allowing the software to choose the strategy to determine best fit. Then we performed a visual inspection of the alignments to check for non big gaps appearing on MSA. The conservative sites were obtained from MAFFT online metrics. To find the best fit evolution model for our MSA, we use Model Finder with default parameters implemented on IQ-TREE 2 (Kalyaanamoorthy et al., 2017). Then we run IQ-TREE 2 with the selected model (GTR+F+G4) to conduct a maximum likelihood (ML) analysis in order to obtain the best phylogenetic tree. Bootstrapping was performed to obtain the consensus from 1000 replicates. We report the consensus likelihood phylogenetic tree with support values of boostrapping above 70% (Log-likelihood of consensus tree −123695.792). The ML analysis was performed four times to ensure the topology of the consensus tree. The results were visualized on Figtree V.1.4.4 (Rambaut et al., 2018) and prepared for image publication on inkscape V1.4.2. Silhouette picture was obtained from Phylopic (https://www.phylopic.org/).

### Orthologues multiple sequence alignment and mapping of conserved structural motifs in GCR1 structure

The three-dimensional structure of the *Arabidopsis thaliana* GCR1 protein was obtained through sequence-based prediction using AlphaFold2 (Jumper et al., 2021). To identify conserved functional motifs, a multiple sequence alignment was performed on orthologous GCR1 sequences retrieved from the ORTHO05D007750 orthology tree of the PLAZA Dicots 5.0 platform (Van Bel et al., 2022). Sequences were filtered by length: those shorter than 250 amino acids or longer than 350 amino acids were excluded, resulting in the removal of 24 sequences from an initial set of 127. The excluded sequences were (by Gene ID): BcaB01g01552 (60 aa), BolC6t35198H (61 aa), SEGI_18851 (87 aa), Lj7A209G42 (90 aa), MSTRG.1710 (101 aa), Lalb_Chr23g0268021 (111 aa), Lalb_Chr23g0268011 (117 aa), SEGI_23264 (118 aa), HPP92_016211 (142 aa), ATR0684G160 (147 aa), MD15G1048200 (165 aa), PGSC0003DMG401012996 (167 aa), BcaB01g01553 (169 aa), unitig_8.g4842 (179 aa), Zm00001eb278600 (209 aa), ClCG05G015850 (375 aa), BolC6t35406H (382 aa), COL.COLO4_34509 (420 aa), SalBow1G5335 (425 aa), MCO03G156 (437 aa), vmacro04115 (442 aa), CDE06G0461 (528 aa), Haze_18321 (643 aa), and PRCOL_00005697 (868 aa).

The alignment was performed using the online T-Coffee server (Notredame et al., 2000) to assess the conservation of key residues across the major evolutionary lineages of *Viridiplantae*. Structural motifs were selected for analysis when their sequential and structural positions corresponded to functional regions described in animal GPCRs, primarily from families A and B (Fredriksson, Lagerström, et al., 2003; Isberg et al., 2015; Venkatakrishnan et al., 2013) These motifs were manually mapped onto the three-dimensional GCR1 model using PyMOL (Schrödinger, LLC).

- **FxxP^5.50^**: Motif present in all GPCR families (Isberg et al., 2015; Taddese et al., 2013)
- **CW^6.50^xP**: Present in family A, associated with receptor activation switches; in GCR1 it appears as SW6.50xF (Hauser et al., 2021; Venkatakrishnan et al., 2013).
- **PxxG^6.48^**: Associated with TM6 disruption upon activation and G alpha coupling in family B; in GCR1 there appears to be a duplication, occurring as PxxxxG^6.48^ (Hilger et al., 2021).
- **YLH^3.50^ / TLH^3.50^**: Motif involved in maintaining the inactive conformation in family B (Kim et al., 2025)
- **TM3–ECL2 disulfide bridge**: Structurally conserved across all families, essential for proper folding of the extracellular loop 2 (Fredriksson, Lagerström, et al., 2003).
- **Sodium ion-binding motif**: In class A GPCRs, centered on residue D2.50 with contributions from S3.39, W6.48, N7.45, and S7.46 (Katritch et al., 2014; White, 2018). In GCR1, D2.50 is preceded by the YLxxxD2.50 motif, characteristic of the family (Taddese et al., 2013), and only residues S3.39 and W6.48 are present.
- **SV^8.50^xxxI in helix 8**: Similar to the EF/V^8.50^xxxL motifs described in families A and B, implicated in H8 stabilization.

## AUTHOR CONTRIBUTIONS

Valentina Fernandez-Figueroa, Formal analysis, Data curation, Investigation, Visualization, Methodology, Writing – original draft.

Camila A. Quercia-Raty, Formal analysis, Data curation, Methodology, Visualization, Writing – review and editing.

Jan Gallastegui-Ulloa, Formal analysis, Data curation, Methodology, Visualization, Writing – review and editing.

Luka Robeson, Formal analysis, Data curation, Methodology, Visualization, Writing – review and editing.

Sebastian E Brauchi, Conceptualization, Data curation, Formal analysis, Methodology, Project administration, Supervision, Writing – original draft, review and editing.

## ACKNOWLEDGEMENT

This research was supported by the Patagon supercomputer of Universidad Austral de Chile (FONDEQUIP EQM180042). We thank CThB for helpful suggestions and improvements to the design of structural figures. No conflict of interest declared.

## Declaration of generative AI and AI-assisted technologies in the writing process

During the preparation of this work, the authors used ChatGPT by OpenAI to correct the language and improve the readability of the text. After using this tool, the authors reviewed and edited the content as needed and took full responsibility for the content of the published article.

